# Design of optimal labeling patterns for optical genome mapping via information theory

**DOI:** 10.1101/2023.05.23.541882

**Authors:** Yevgeni Nogin, Daniella Bar-Lev, Dganit Hanania, Tahir Detinis Zur, Yuval Ebenstein, Eitan Yaakobi, Nir Weinberger, Yoav Shechtman

## Abstract

Optical genome mapping (OGM) is a technique that extracts partial genomic information from optically imaged and linearized DNA fragments containing fluorescently labeled short sequence patterns. This information can be used for various genomic analyses and applications, such as the detection of structural variations and copy-number variations, epigenomic profiling, and microbial species identification. Currently, the choice of labeled patterns is based on the available bio-chemical methods, and is not necessarily optimized for the application. In this work, we develop a model of OGM based on information theory, which enables the design of optimal labeling patterns for specific applications and target organism genomes. We validated the model through experimental OGM on human DNA and simulations on bacterial DNA. Our model predicts up to 10-fold improved accuracy by optimal choice of labeling patterns, which may guide future development of OGM bio-chemical labeling methods and significantly improve its accuracy and yield for applications such as epigenomic profiling and cultivation-free pathogen identification in clinical samples.

## 1 Introduction

Optical genome mapping (OGM) (Deen, Vranken, et al., 2017; Jeffet et al., 2021; Levy-Sakin and Ebenstein, 2013; Neely, Dedecker, et al., 2010) is a technique for mapping optically imaged linearized DNA fragments to reference genome sequences. It has been demonstrated for the detection of structural variations and copy-number variations (Ebert et al., 2021), DNA damage (Torchinsky et al., 2019), epigenomic profiling (Gabrieli, Michaeli, et al., 2022; Gabrieli, Sharim, et al., 2018), and microbial species identification (Bouwens et al., 2020; Grunwald et al., 2015; Müller, Nyblom, et al., 2020; Wand et al., 2019). The current common process by which OGM is implemented involves linearizing (or combing, stretching) the labeled DNA fragments on some surface (Deen, Sempels, et al., 2015; Levy-Sakin and Ebenstein, 2013; Wu et al., 2018), followed by optical measurement of the density of a specific short genome sequence pattern along the DNA fragments, by the fluorescent labeling of the pattern occurrences (Neely, Dedecker, et al., 2010), and the analysis of the acquired images. The mapping of the images to reference genome sequences is done using alignment algorithms (Bouwens et al., 2020; Dehkordi, Luebeck, and Bafna, 2021; Mendelowitz and Pop, 2014; Valouev et al., 2006).

Although OGM extracts only partial information from the genome, it possesses several advantages compared to DNA sequencing. These include the ability to generate extremely long reads of up to megabase size, which is necessary for detecting genomic large-scale structural and copy number variations, and the potential for extremely high sensitivity, i.e. detection of low quantities of target DNA (Margalit et al., 2021), which is necessary in applications such as cultivation-free pathogen identification (Müller, Nyblom, et al., 2020; Nyblom et al., 2023).

Multiple labeling methods are used in OGM. Originally, restriction enzymes were used and the visible cut sites were used as labels (Schwartz et al., 1993). More recently, fluorescent labeling is done to label short sequence patterns (Neely, Dedecker, et al., 2010). One labeling approach is competitive binding based (Müller, Dvirnas, et al., 2019; Müller, Nyblom, et al., 2020; Nyblom et al., 2023), where a fluorophore competes with a DNA intercolating molecule for binding to the DNA, allowing the visualization of G/C base-pair density along a DNA fragment. Another approach uses CRISPR-Cas9 DNA labeling with a multitude of sgRNAs (Abid et al., 2020). One more labeling approach is achieved by enzymes from bacterial restriction-modification (RM) systems that have short recognition sequences of 4-6 base-pair (bp) length, such as nicking endonucleases, with incorporation of fluorescently labeled nucleotides (Neely, Deen, and Hofkens, 2011), and DNA methyltransferases (DNMT) that methylate a specific sequence pattern, and together with modified co-factors, their action is modified to fluorescent labeling (Dalhoff et al., 2006; Deen, Vranken, et al., 2017; Grunwald et al., 2015; Pljevaljčić, Schmidt, and Weinhold, 2004). The REBASE database (Roberts et al., 2015) lists thousands of restriction-modification systems, with different recognition sequences, many of which are available commercially. This suggests a wide playground for optimization of the labeling patterns for specific applications and target genomes, given a way to quantify and predict OGM accuracy for each pattern.

Previous attempts to computationally estimate the accuracy of OGM were done for example in (Bouwens et al., 2020; Wand et al., 2019), where simulated estimations of OGM accuracy were done, mainly tailored to the specific labeled patterns and algorithms used in those studies. Before the advent of fluorescent labeling based OGM, in (Anantharaman and Mishra, 2001), p-value estimation of restriction map alignment was done using combinatorial methods. In this paper, we propose an information theory (Cover and Thomas, 2012) based method for the labeled pattern design. Information-theoretic analysis has been applied for various DNA processing problems. For example, (A. S. Motahari, Bresler, and Tse, 2013) analyzed the minimal read coverage needed for genome reconstruction DNA shotgun sequencing, mostly in a noiseless setting. Information-theoretic analysis of reference-based DNA shotgun sequencing has been derived in (Mohajer, A. Motahari, and Tse, 2013), and recently refined in (Weinberger and Shomorony, 2023). Information-theoretic methods were also applied to the analysis of nanopore DNA sequencing (Mao, Diggavi, and Kannan, 2018), where the nanopore channel was modeled as an insertion-deletion channel and general bounds on the number of possible decodable sequences were developed. However, tight bounds on the error probability were not developed.

In this work, we develop a model of OGM based on information theory and compute its error probability, thus enabling the optimization of labeling patterns. To achieve this goal, we model OGM as a random code operating over a noisy communication channel (Shannon, 1948) and use tight bounds on its error probability. We present experimental and simulated validations of the model, and demonstrate the optimization of labeling patterns for different target genome sets, such as the human genome and bacterial genomes. This paper is organized as follows. Section 2 develops the information-theoretic based model of OGM and describes the simulation and experimental methods we used for its validation. Section 3 presents the results of the validation for Error probability vs. DNA fragment length, and the results of the application of the model to the design of optimal labeling patterns.

## 2 Materials and Methods

In OGM, a specific short genome sequence pattern is labeled in observed DNA fragments (Figure 1). Given a set of target genome sequences from which the observed DNA fragments originate, and a specific labeled sequence pattern, we are interested in the accuracy (or error probability) for mapping each fragment to its true position from one of the target genome sequences. The target genome sequences may consist, for example, of the human genome, a set of bacterial genomes, etc.

**Figure 1:**
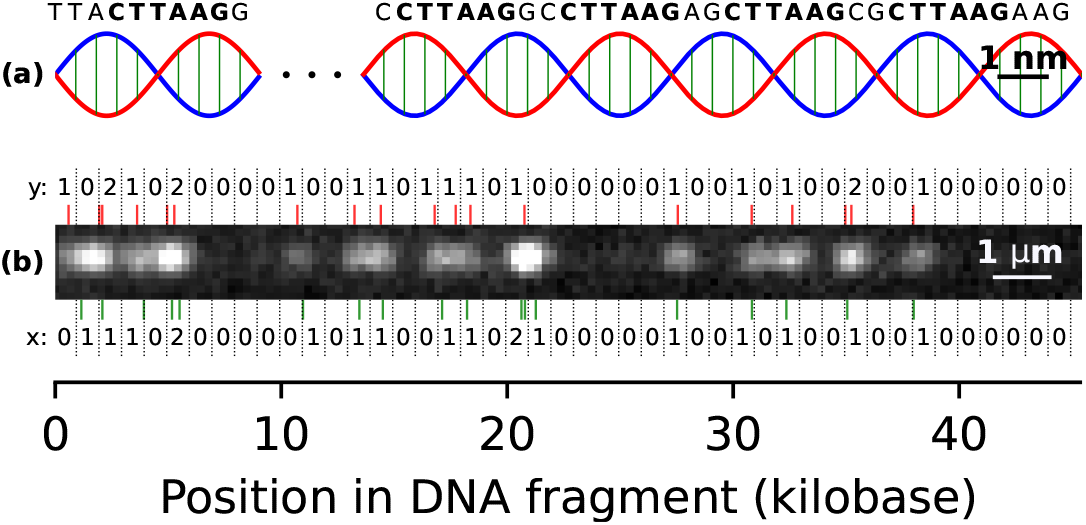
PatternCode concept. Subplot (a) presents an illustration of a DNA molecule (not to scale), with the pattern CTTAAG labeled. Subplot (b) shows an example DNA fragment experimental image, which is used for estimation of theoretical model parameters. Bottom: the pattern occurrences in the human genome sequence, matching the DNA fragment, along with their bin counts (*x*). Top: localized labels, together with their bin counts (*y*).

A full model of the OGM process would include bio-chemical labeling errors of DNA fragments, non-uniform stretching of DNA fragments, optical imaging artifacts, and localization errors of labels in the images. Here, the model is simplified to enable theoretical analysis; nevertheless, our experimental validations, detailed below, show that it constitutes an excellent approximation of the actual measurement process.

### 2.1 Information theory of OGM accuracy

To simplify the model and estimate the error probability, both the genome sequence and DNA fragments (Figure 1) are segmented into non-overlapping bins of size *B*, typically on the order of 1 kilobase (kb). The chosen value of *B* is explained in Supplementary Figure S1. When the bin size is too small, label localization errors cause statistical dependence between bins; and if it is too large, significant information regarding the label position is lost. Here, statistical independence of the information contained in neighboring bins is assumed, allowing for the modelling of the OGM process as a discrete memoryless channel (DMC) decoding problem in the random coding regime (Gallager, 1968; Shannon, 1948).

We model the OGM process as a noisy communication channel, transmitting a message (DNA fragment position in the genome) which is encoded by a codeword (the bin counts corresponding to pattern positions in the sequence), resulting in an information-theoretic code we term the pattern-code. At the output of the communication channel a noisy codeword (image of the DNA fragment) is received, and is decoded to produce the decoded message (estimated genome position), out of the possible messages, namely the codebook (all possible genome fragments). The process is summarized in Figure 2. The codebook of size *M* is composed of codewords which are vectors of bin counts of length *n*. Each codeword is associated to a message of position offset, and corresponds to a segment of a genome sequence of length *L* = *nB*, where *B* is the bin size. Each value *x* in the codeword is the number of pattern occurrences in a bin of the genome sequence segment. Defining *G* as the total genomes length (sum of lengths of all target genome sequences), the size of the codebook is *M* = 2*G/B* (approximately, considering valid position offsets), where the factor of 2 is due to the unknown orientation of the DNA fragment, requiring consideration of both the forward and reverse direction of the fragment. The measured noisy codewords are the imaged DNA fragments, and are vectors of bin counts of length *n*. Each value *y* in the vector is the number of detected labels in a bin of the DNA fragment. The channel model is described by the likelihood *p*_*y*|*x*_, which captures the various noise factors in the OGM process. The mapping algorithm of OGM, in this case, is a maximum likelihood (ML) decoder which chooses the codeword (and message) that maximizes the likelihood of the observed noisy codeword.

**Figure 2:**
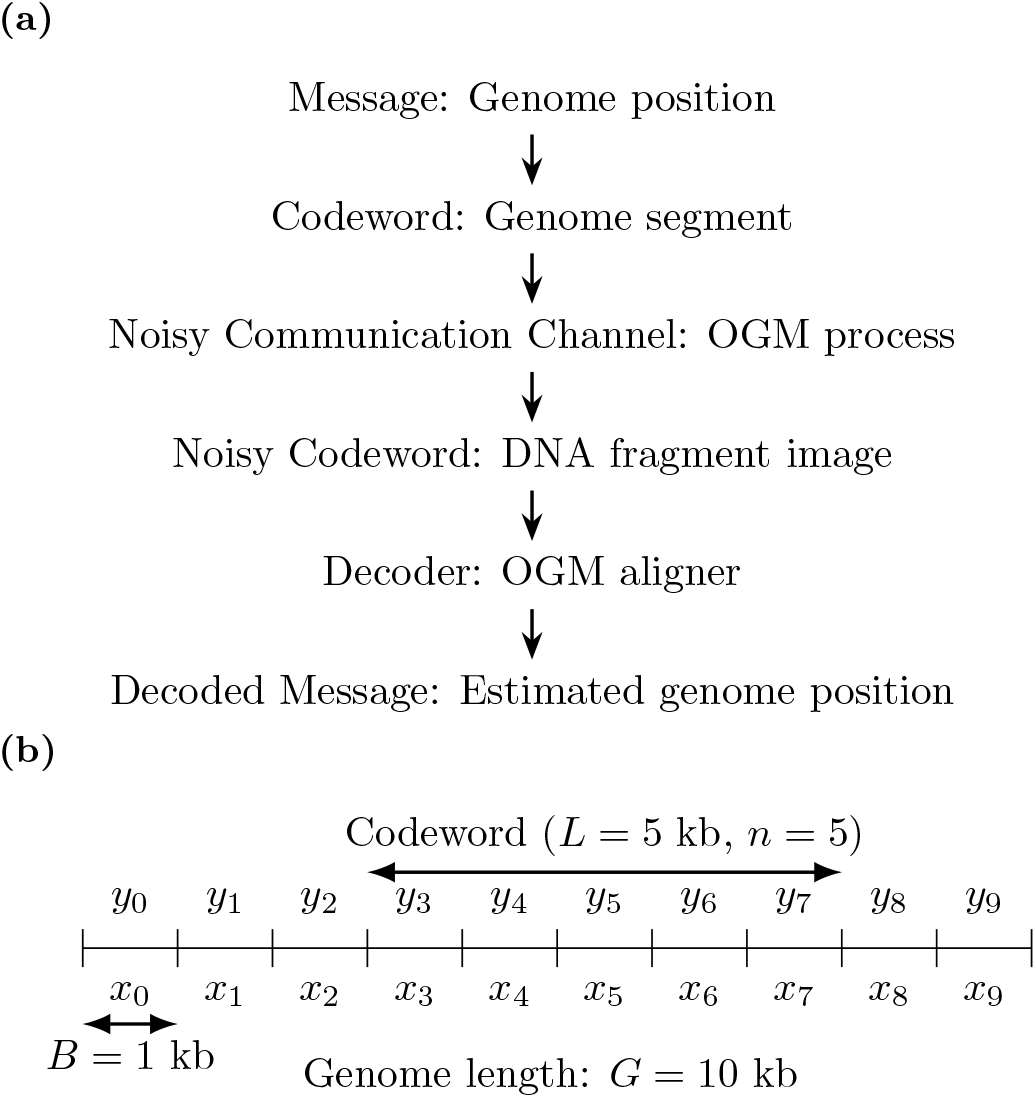
Information-theoretic model of OGM as a random codeword transmitted over a noisy communication channel. (a) The OGM communication channel flow chart. (b) Example model parameters, explained in Section 2.1. *B* is the bin size, *L* is the DNA fragment length, *n* is the number of bins in the DNA fragment, and *G* is the genome length (here it’s very short, for illustration). The number of possible position offsets of a codeword in the binned genome is the number of messages (or codebook size) *M*. Also shown are the bin counts of pattern occurrences in the genome sequence (*x*) and of the labels in the DNA fragment image (*y*).

Modeling the problem in this way allows to use sharp and accurate bounds from the information-theoretic literature to bound the error probability in OGM. The celebrated noisy channel coding theorem (Shannon, 1948) assures that optimal maximum-likelihood decoder achieve a vanishing probability when the coding rate is less than the *capacity* of the channel. Importantly, this is proved using a randomly chosen codebook, for which the codeword symbols are randomly chosen. This vanishing error probability is obtained for asymptotically large codeword size *n* → ∞. However, in the case of OGM, we are interested in the error probability for the shortest possible DNA fragments, and relatively high error probabilities can often be tolerated, as final clinically relevant decisions can be based on multiple mapped DNA fragments, allowing for the correction of errors. In recent years, tight bounds and approximations on the coding rate for non-vanishing error probabilities in DMCs has been obtained (Hayashi, 2009; Polyanskiy, 2010; Polyanskiy, Poor, and Verdú, 2010; Tan et al., 2014), and these are the bounds we will henceforth utilize.

#### 2.1.1 Error probability computation

Having formulated the OGM problem as the transmission of a codeword from a random codebook over a DMC, we can compute the error probability of the maximum-likelihood decoder following the achievability and converse probability bounds for a DMC in (Tan et al., 2014, Equations 4.33, 4.57). The error probability, denoted by *ε*, is approximated as follows:

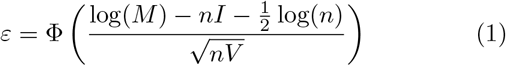

where F is the standard normal cumulative distribution function, *M* = 2*G/B* with *G* the total genomes length (sum of lengths of all target genome sequences), *n* = *L/B* with *L* the DNA fragment length, and *B* the bin size as above. The mutual information *I* (of *x* and *y*) and the variance *V* are computed from the joint and marginal distributions of the number of pattern occurrences in a bin of a genome sequence (*x*) and the number of detected labels in a bin of a DNA fragment (*y*), as follows. We define *r* as the following log-ratio of joint and marginal distributions:

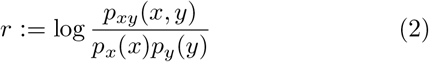

where *p*_*xy*_ = *p*_*y*|*x*_*p*_*x*_ is the joint distribution of *x* and *y*, and *p*_*y*_ = E_*x*_[*p*_*y*|*x*_] is the marginal distribution of *y*. Then, *I* = E_*xy*_[*r*] and *V* = Var_*xy*_[*r*]. The only free parameters in this model that need to be estimated from the data are the pattern density distribution *p*_*x*_ and the label detection likelihood *p*_*y*|*x*_.

#### 2.1.2 Estimation of parameters

The pattern density distribution *p*_*x*_ was estimated from the target genome sequences by computing the histogram of *x*, the number of pattern occurrences in all the bins of size *B*. The label detection likelihood *p*_*y*|*x*_ was estimated from experimental OGM data by computing the 2-d histogram of (*x, y*) pairs of aligned bins over all DNA fragments (see Table 1). Here, *x* is the number of pattern occurrences in a bin of the target genome sequence, and *y* is the number of detected labels in a bin of a DNA fragment.

**Table 1:**
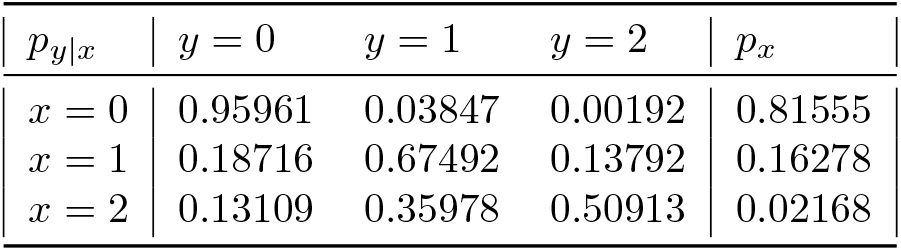
Estimated parameters of information-theoretic noisy DMC model used in this work. The pattern density distribution *p*_*x*_ for the pattern CTTAAG is shown on the human genome sequences. The label detection likelihood *p*_*y*|*x*_ was estimated from experimental human DNA fragments, and the human genome hg38 with the same pattern labeled, as described in Section 2.1.2. The number of pattern occurrences in a genome bin, *x*, was capped at 2, and the number of labels detected in a DNA fragment bin, *y*, was capped at 2. The bin size used was 1*kb*.

The likelihood used in this work was estimated for experimental human DNA fragments, and the human genome hg38 (Figure 1), where the pattern CTTAAG was labeled. This data consisted of images acquired as described in Section 2.4, where the DNA fragments were labeled by the DLE-1 enzyme and imaged using the Saphyr system (Bionano Genomics). The DNA fragments, all longer than 400kb, were aligned to the human genome, using the DeepOM algorithm (Nogin et al., 2023). Overall, 445 DNA fragments were used consisting of a total of 180 megabase (Mb).

In both estimated probabilities, and throughout this work, the *x* and *y* counts were capped at the number 2 (i.e. if more than 2 pattern occurrences or more than 2 labels were detected in a bin, the count was set to 2). This can be justified by the low density of length-6 patterns in the genome (see *p*_*x*_(*x* = 2) in the table) and the inability of practical localization methods to separate closely spaced labels.

### 2.2 Simulated validation of the theory

The simulation-based validation of the theory in Equation (1) was done by simulating DNA fragments of length *L* = *nB* bp, each at a random start position in target genome sequences. The counts of labels in each bin (*y*) were simulated from the *p*_*y*|*x*_ distribution, where *x* is the number of pattern occurrences in a bin of a genome sequence and *y* is the number of detected labels in a bin of a DNA fragment (see Section 2.1). The maximum-likelihood decoder estimates the position offset *î* of a DNA fragment in a genome sequence as follows:

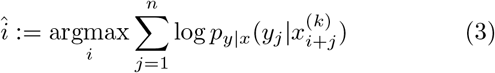

where 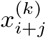 is the bin-count of pattern occurrences in the *j*-th bin of the genome sequence *k* starting at the bin from position *i*. For each genome sequence, its reverse sequence was also considered, since the orientation of the fragment is unknown. The error rate was computed by running the decoder on all generated DNA fragments (512 per parameter set: over a multitude of fragment lengths, labeled patterns, genome sets, see Figures (3,4)), with a decoder error defined when the estimated position is different from the true position.

**Figure 3:**
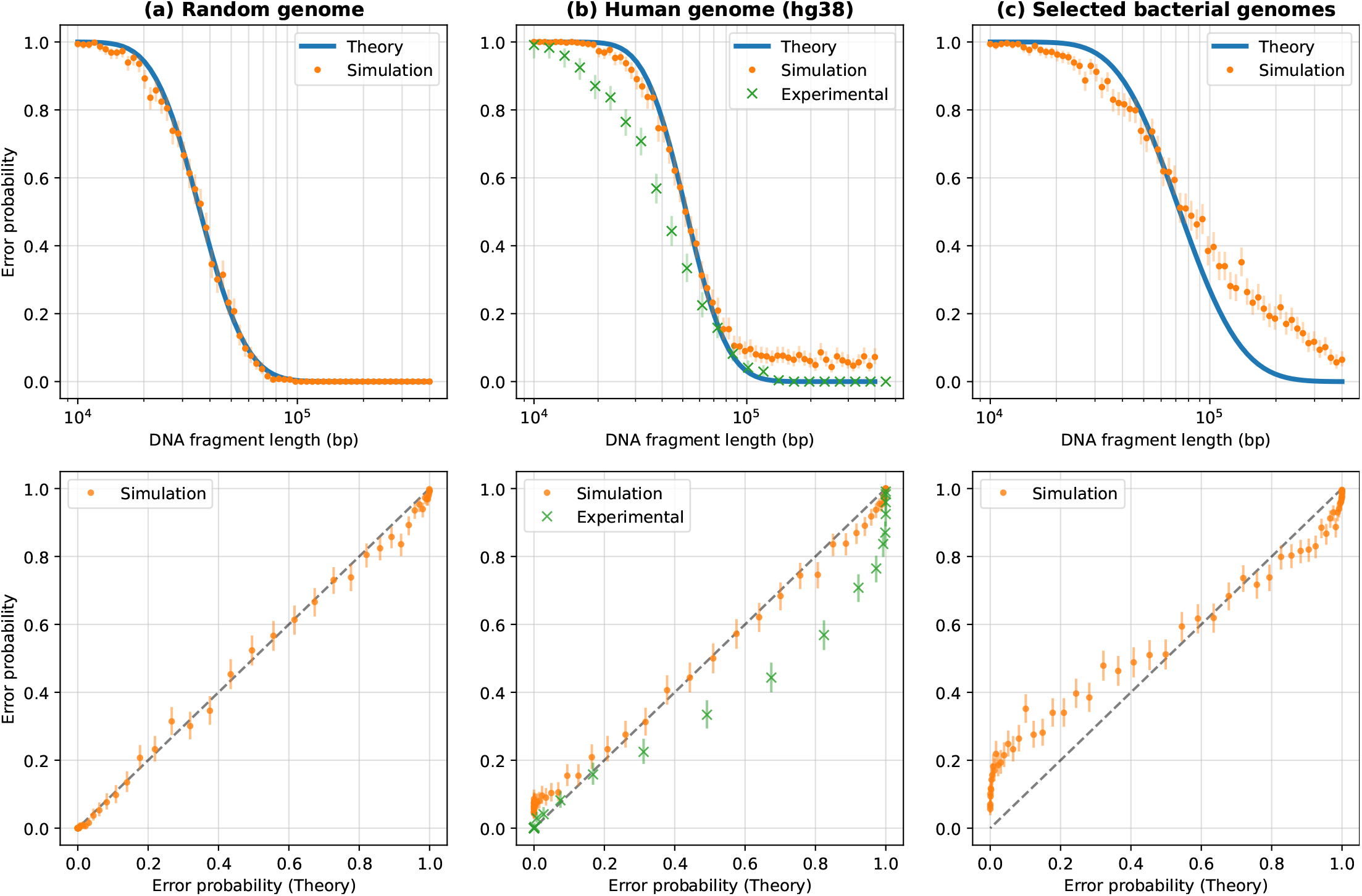
Validation of the theoretical model vs. DNA fragment length. Validation results are shown against three sets of genome sequences: a random genome sequence of length 10^8^ bp (subplot a), the human genome (subplot b), and selected bacterial genomes (subplot c; see Table S1). The plots in the top row show the error probability versus fragment length, while the bottom row plots show the same data but with the theoretical error probability on the x-axis and the simulated or experimental error rates on the y-axis. The theoretical error probability was computed using Equation (1), and the simulated error rate was computed as described in Section 2.2, based on 512 randomly generated DNA fragments for each length shown. For the human genome, in addition to the simulated error rates, we show the experimental error rates for comparison. These experimental error rates were adapted from Figure 4b in (Nogin et al., 2023) and described in Section 2.3. For both the simulated and experimental error rates, 95% confidence bounds (Clopper and Pearson, 1934) for the rate are shown, based on the error counts and the number of trials.

**Figure 4:**
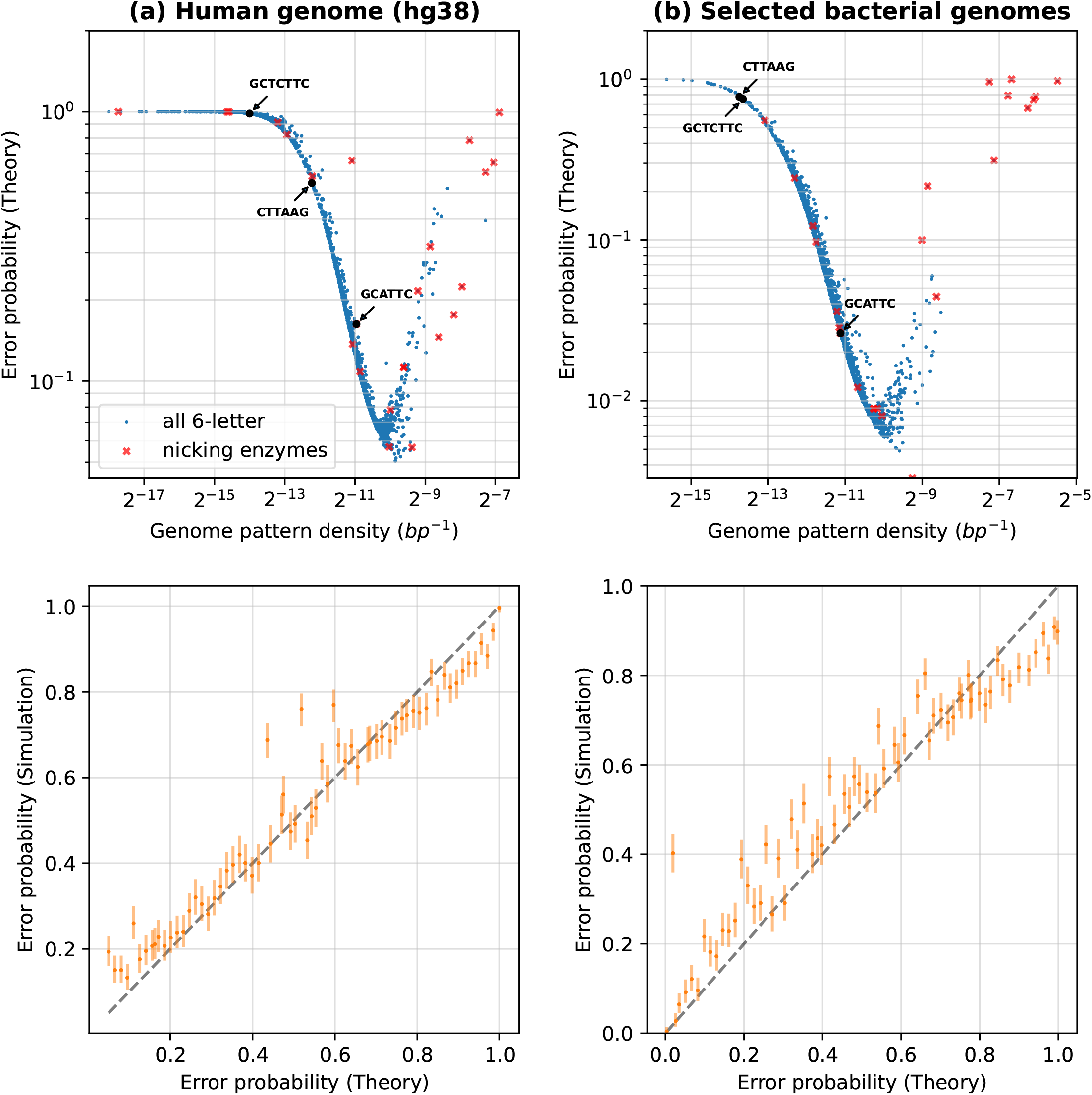
Design of optimal labeling patterns. This figure shows the evaluation of error probabilities for all length-6 labeling patterns, and some special patterns, against the following sets of genome sequences for a fixed DNA fragment length: subplot (a) is for the human genome, subplot (b) is for selected bacterial genomes (see Table S1). We varied the labeling pattern, with all 4^6^ = 4096 possible patterns evaluated and shown on the subplots. Additionaly, highlighted are nicking enzyme recognition patterns from the REBASE database and some commonly used enzyme recognition patterns in OGM are shown with arrows (DLE-1, Nt.BspQI, Nb.BsmI with recognition patterns CTTAAG, GCTCTTC, and GCATTC respectively). For each pattern, its reverse complement is also labeled at the same time. The data plotted for these special patterns for the human genome is shown in Supplementary Table S2. For each genome set, we computed the theoretical error probability using the estimated *p*_*x*_ and *p*_*y*|*x*_ (see Section 2.1), and validated the theory through simulations (see Section 2.2). The top row of subplots shows the error probability vs. the density of the labeling pattern, while the bottom row plots compare the theory (x-axis) with simulated evaluation (y-axis) of the error probability for a random subset of patterns, with 95% confidence intervals computed as in Figure 3.

### 2.3 Experimental validation of the theory

The experimental validation of the theory in Equation (1) was done by comparing it to results from the DeepOM work (Nogin et al., 2023), which evaluated the error rates of the DeepOM OGM algorithm in mapping DNA fragments to the genome, using experimental human DNA fragments labeled with the pattern CTTAAG (Figure 1). The algorithm is based on a Convolutional Neural Network (CNN) for localizing labels in a DNA fragment image, and a Dynamic Programming (DP) algorithm for aligning the DNA fragment to the genome. Overall, 512 DNA fragments were used (per fragment length, over multitude of fragment lengths) consisting of a total of 1.43 gigabase (Gb).

### 2.4 Sample preparation and data collection

Images of human DNA fragments (a total of 445 images, see one example in Figure 1), were used to estimate the label detection likelihood parameters (Section 2.1.2). These images were captured using the Saphyr instrument and Saphyr chips (G1.2) (Bionano Genomics). The protocol for extraction and labeling of DNA is described in Nogin et al. (Nogin et al., 2023). U2OS human cell line was cultured in Dulbecco’s Modified Eagle medium with 10% fetal bovine serum (Gibco, Amarillo, TX), 2 mM l-glutamine, and 1% penicillin-streptomycin (10,000 U/mL; Gibco). The cells were incubated at 37°C with 5% CO_2_. DNA was extracted from 10^6^ cells using the Bionano Prep Cell Culture DNA Isolation Protocol (Bionano Genomics). 1 μg of DNA was directly labeled using the Bionano Genomics DLS labeling kit, composed of a single enzymatic labeling reaction with DLE-1 enzyme. The reaction mixture contained 6 μl of 5x DLE-buffer, 2 μl of 20x DL-Green, 2 μl of DLE-1 enzyme (Bionano Genomics), and a total reaction volume of 30 μl. The reaction was incubated for 2 hours at 37°C.

### 2.5 Genome sequence data

Human and bacterial genome sequence data was downloaded from the NCBI genome datasets https://www.ncbi.nlm.nih.gov/datasets/. Human genome release GRCh38.p14 was used. The bacterial genomes used are listed in Table S1.

## 3 Results

### 3.1 Accuracy vs. fragment length

We validate our theoretical model by comparing it to experimental and simulated data for varying DNA fragment lengths. We first estimate the label detection likelihood *p*_*y*|*x*_ from the experimental data, as described in Section 2.1.2, resulting in the values shown in Table 1. We then assume that the label detection noise is the same for all genome sets, as it is a function of the imaging instrument and the labeling protocol. For each genome set, we estimate the pattern density distribution *p*_*x*_ as described in Section 2.1.2. We then compute the theoretical error probability as described in Section 2.1, using the estimated *p*_*x*_ and *p*_*y*|*x*_. We validate the theory through simulations as described in Section 2.2. For the human genome, we also compare to the experimental data from DeepOM (Nogin et al., 2023).

The results are shown in Figure 3, which shows the error probability vs DNA fragment length. For the random genome sequence, there is an almost perfect match between the theoretical and simulated error probabilities, validating the information-theoretic model in Section 2.1. For both the human and bacterial genomes, there is remarkable match between the theoretical and simulated error probabilities for shorter DNA fragment lengths, meaning that, to a good approximation, real genomes behave as random sequences, with respect to the distribution of labeled patterns. For longer fragments, the simulated probability is higher than the theoretical, which can be explained by the fact that the theoretical model assumes that the labeling pattern is uniformly distributed in the genome, while in reality, genome sequences contain a multitude of non-uniformly distributed anomalous regions, such as repeated sequences, unmapped regions in the centromeres and telomeres in the human genome, highly similar regions in closely related bacterial species, etc. For the human genome, the experimental error rate (see Section 2.3) is slightly lower than the theoretical prediction. This can be explained by the fact that the DeepOM alignment algorithm uses the exact localized positions of the labels for the alignment to the genome sequence, which is added information that is not accounted for in the theoretical model, and aids in the accuracy of the alignment. Also, the experimental error rate vanishes for long enough fragments, which doesn’t happen in the simulation, due to the fact that the experimental error rate in the respective study was computed only for fragments from relatively conventional and non-anomalous regions of the genome. With all that said, the theoretical model still provides a good upper bound on the achievable error rate.

### 3.2 Design of optimal labeling patterns

Now that we have validated the theory for the accuracy of OGM for a specific labeled pattern (CTTAAG) on the human genome using experimental and simulated data, we can go beyond analysis to synthesis: the design of optimal labeling patterns, which is a direct practical implication of this work.

The results are shown In Figure 4, where we show the error probability for different labeling patterns. The DNA fragment length was fixed, and the labeling pattern was varied. For a subset of patterns, we validated the theory by simulation as described in Section 2.2.

The examined patterns were generated as follows: all 4^6^ = 4096 possible sequences over *A, C, G, T* of length 6 were generated, with the pattern positions chosen as the union of its positions in the genome and its reverse complement (Section 2). Note that for palindromic patterns (there are 4^3^ = 64 such patterns), the reverse complement is identical to the pattern itself. Additionally, some special patterns of interest were examined (see Supplementary Table S2): nicking enzyme recognition patterns (with IUPAC codes expanded to matching sequences) from the REBASE database (Roberts et al., 2015), and some commonly used enzymes in OGM. For each pattern, we computed its average density in the genome by dividing the number of pattern occurrences by the genome length. We assumed for all patterns the same label detection likelihood *p*_*y*|*x*_ as used in Section 3.1. This is valid under the assumption that other bio-chemical labeling methods that have 6-letter recognition sequences would have similar labeling errors.

It can be seen in the figure, that the error probability has strong dependence on the pattern density, achieving a minimum for specific patterns with a density on the order of 1 kilobase (kb). While for a random genome sequence, one would expect the length-6 pattern density to be on the order of 4^*−*6^ (Supplementary Figure S2), real genome sequences behave differently, because the labeled sequences are not uniformly distributed (Figure 4). For example, some 6-letter sequences are much more frequent in the genome than others. The figure shows that for both the human genome and the bacterial genomes, the best patterns have a predicted error probability which is more than 10-fold lower than the commercially available pattern we examined experimentally, suggesting there is significant room for improvement. For example, as a direct prediction from our model, the best 6-letter sequence for OGM of the human genome, would be GGAGGC (see supplementary data table file human genome p err vs pattern.csv for the results for all patterns), which would lead to 10X improved accuracy, directly translatable to experimental parameters such as acquisition speed, low sample DNA quantity, simplified sample preparation, etc. In general, patterns with low density are less informative because there are too few labels on the DNA fragment to map it accurately to the reference genome, whereas patterns with high density patterns are less informative because the label detection error per bin is too high. This leads to the conclusion that for a given target genome set with pattern density distribution *p*_*x*_ and label detection likelihood *p*_*y*|*x*_ of the labeling and imaging system, the theory can predict specific optimal labeling patterns which can be more than an order of magnitude better in terms of accuracy.

## 4 Discussion

We showed that our information-theoretic model provides a good approximation for the error probability of OGM. It depends on only four parameters: the target genome length, the DNA fragment length, and two easily estimated parameters: the label detection likelihood, estimated from experimental genome-aligned DNA fragment images, and the labeling pattern distribution, estimated from the target genome sequences. This enables the design of better OGM experiments without the assumption of a specific OGM algorithm, and allows for the intuitive understanding of the importance of different parameters on the accuracy, such as the logarithmic dependence on the target genome length versus the polynomial dependence on the fragment length Equation (1). Additionally, the model enables fast computation due to its simple analytical form, allowing for the design of protocols where multiple patterns are labeled with multiple labeling reagents through combinatorial optimization of pattern combination selection.

In the model, we neglected several factors that might affect the actual error probability in real OGM experiments. The model assumes that an OGM algorithm has access only to bin-counts of labels (Section 2.1), while there is more information that can be utilized from localized positions of labels within each bin. Modeling the process as a DMC, statistical independence of the information in neighboring bins was assumed, which was mitigated by choosing a bin size that is larger than the label localization error margin (Figure S1). In terms of the codebook model, due to overlaps and unknown orientation of the codewords, this implies that the codewords are not exactly independent and identically distributed (as required for the validity of the noisy channel coding theorem). The random codebook model is still rather accurate since a codeword is independent from almost all the other codewords in the codebook. We also assumed that the bin start positions are aligned with the reference genome, while in practice, the bin start positions are not known, so the reference genome sequence can be shifted by a random amount from the true position, up to the bin size. We also neglected the effect of non-uniform stretching of the DNA fragments, before they are imaged, thus the assumption of one-to-one mapping between consecutive bins in the reference genome and the DNA fragment image is not exact and will increase the size of the effective codebook. However, the dependence on the codebook size is logarithmic in Equation (1), so this effect is secondary in comparison to pattern density and fragment length.

Additionally, we applied our model to the design of optimal OGM labeling patterns for the human genome and selected bacterial genomes. However, we made some underlying assumptions that might impact the validity of the results. The label detection likelihood *p*_*y*|*x*_ (Table 1) was estimated (Section 2.1.2) for a specific labeling protocol and reagent, and might be different for other labeling reagents used to label other patterns. We limited our analysis to length-6 patterns, as this is the length of the pattern we examined experimentally. However, the theory can be applied to other pattern lengths, given the label detection likelihood *p*_*y*|*x*_ is estimated from relevant experimental data. It is important to note that some reagents, which can be used for the labeling processes, have recognition sequence patterns with characters matching more than one base type. This means that the labeling pattern is a set of recognition sequences, which we did not consider in this analysis, but it is straightforward to do so.

To conclude, we developed an information-theoretic model of OGM, which enables the prediction of its accuracy and the design of optimal labeling patterns for specific applications and target organism genomes. The accuracy is predicted using parameters easily estimated from target genome sequences and experimental data. The model was validated experimentally on human DNA with a specific pattern labeled, and through simulations with varying DNA fragment lengths, various labeling patterns, and various genomes, including the human genome, bacterial genomes, and randomly generated genome sequences. The model predicts up to 10-fold better accuracy by optimal choice of labeling pattern for the human genome and bacterial genomes. Future development of bio-chemical reagents and protocols for labeling the patterns suggested by the model may significantly improve the accuracy and yield of OGM for applications such as epigenomic profiling and cultivation-free pathogen identification in clinical samples.

## Data Availability

All code and data used to generate the results of this work is deposited on Zenodo and is publicly available at DOI: https://doi.org/10.5281/zenodo.7957345. The numerical results of error probabilities for each pattern in Figure 4 and Supplementary Figure S2 are attached as supplementary csv files:

- `Human Genome:` `human_genome_p_err_vs_pattern.csv`
- `Bacterial Genomes:` `bacterial_genomes_p_err_vs_pattern.csv`
- `Random Genome:` `random_genomes_p_err_vs_pattern.csv`

