## Supplementary Information for "Design of optimal labeling patterns for optical genome mapping via information theory"

May 22, 2023

| | Organism | Accession | Sequence ID | Sequence Length |
|---|---|---|---|---|
| 0 | Citrobacter koseri ATCC BAA-895 | GCF_000018045.1 | NC_009792.1 | 4720462 |
| 1 | Escherichia coli str. K-12 substr. MG1655 | GCF_000005845.2 | NC_000913.3 | 4641652 |
| 2 | Klebsiella pneumoniae subsp. pneumoniae HS11286 | GCF_000240185.1 | NC_016845.1 | 5333942 |
| 3 | Mycobacterium tuberculosis H37Rv | GCF_000195955.2 | NC_000962.3 | 4411532 |
| 4 | Proteus mirabilis HI4320 | GCF_000069965.1 | NC_010554.1 | 4063606 |
| 5 | Pseudomonas aeruginosa PAO1 | GCF_000006765.1 | NC_002516.2 | 6264404 |
| 6 | Salmonella enterica subsp. enterica serovar Ty... | GCF_000006945.2 | NC_003197.2 | 4857450 |
| 7 | Staphylococcus aureus subsp. aureus NCTC 8325 | GCF_000013425.1 | NC_007795.1 | 2821361 |

Table S1: Selected bacterial species used in this work. These were selected as they are common in clinical samples and are known to be pathogenic.

| | Pattern | Error probability | Genome pattern density ($bp^{-1}$) |
|---|---|---|---|
| 0 | CCGG | 0.056730 | 0.001501 |
| 1 | GCAGTG | 0.056941 | 0.000960 |
| 2 | GGATC | 0.078198 | 0.000983 |
| 3 | GGTCTC | 0.108176 | 0.000538 |
| 4 | GACTC | 0.112301 | 0.001273 |
| 5 | GAGTC | 0.112305 | 0.001273 |
| 6 | GCAATG | 0.136921 | 0.000465 |
| 7 | GASTC | 0.145349 | 0.002546 |
| 8 | GAATGC | 0.162584 | 0.000502 |
| 9 | GCATTC | 0.162584 | 0.000502 |
| 10 | GCWGC | 0.176059 | 0.003426 |
| 11 | GGATG | 0.216034 | 0.001684 |
| 12 | GCNGC | 0.223953 | 0.004022 |
| 13 | GTCTC | 0.315816 | 0.002147 |
| 14 | CTTAAG | 0.543502 | 0.000208 |
| 15 | CGTCTC | 0.576196 | 0.000209 |
| 16 | CCWGG | 0.596975 | 0.006383 |
| 17 | CCNGG | 0.647643 | 0.007523 |
| 18 | CGCG | 0.657824 | 0.000460 |
| 19 | GATC | 0.784620 | 0.004662 |
| 20 | CACGAG | 0.822194 | 0.000127 |
| 21 | RCCGGY | 0.913523 | 0.000107 |
| 22 | GCTCTTC | 0.984998 | 0.000061 |
| 23 | AGCT | 0.994101 | 0.008437 |
| 24 | CTCGAG | 0.998656 | 0.000039 |
| 25 | GCCGGC | 0.999033 | 0.000041 |
| 26 | CGATCG | 1.000000 | 0.000005 |

Table S2: Special patterns used in the data underlying Figure 4. Error probabilities are given for the human genome, for the following selected patterns: Nicking enzyme recognition sequences (4 letters or longer) from (`http://rebase.neb.com/rebase/azlist.nick.html`), and commonly used patterns in OGM (the enzymes DLE-1, Nt.BspQI, Nb.BsmI with recognition patterns CTTAAG, GCTCTTC, and GCATTC respectively)

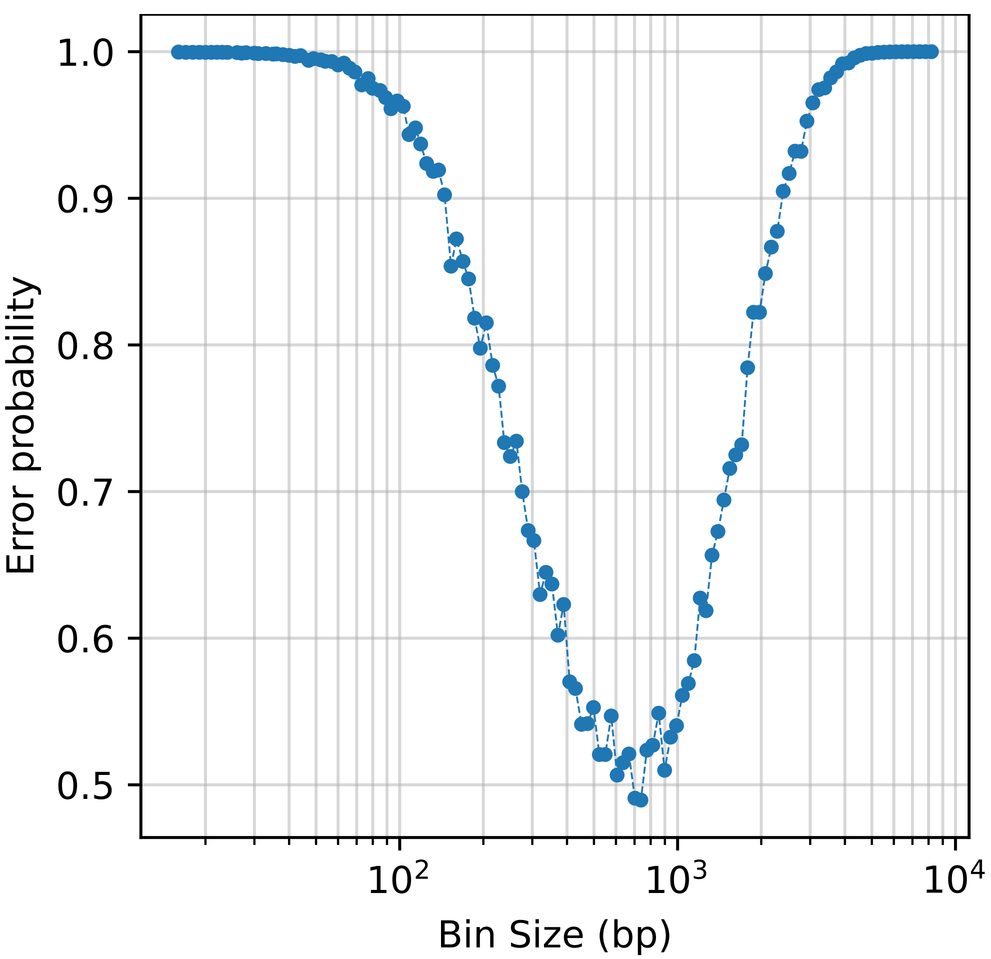


Figure S1: Selection of optimal bin size. To enable the use of the DMC model, the bin size should be larger than the label localization error margin of the localization microscopy method used. If this is not the case, the information in neighboring bins is not independent. To make the assumption of DMC as valid as possible, we optimize the predicted theoretical error probability by varying the bin size. This ensures the tightest possible upper bound on the error probability compared to a decoder (or OGM aligner) that is not limited to binning the labels, the genome sequences, and which does not assume a DMC. The computation of the error probability is done with the same parameters as in Figure 3, for the human genome (DNA fragment length of 50kb, labeled at the pattern CTTAAG). Except that for each bin size, the labeling detection likelihood $p_{y|x}$, as well as $p_x$, are computed for the specific bin size (Table 1, Section 2.1.2). When the bin size is too low, the localization error of labels relative to the pattern position makes the bins dependent statistically, increasing the label count estimation error and the error probability. When the bin size is too large, the DMC assumption is valid, but too much information is lost, as the number of bins (or codeword length $n$) is reduced, and the error probability increases. The plot shows that the optimal value is around 1kb, which is the value we used for the bin size $B$ in this work.

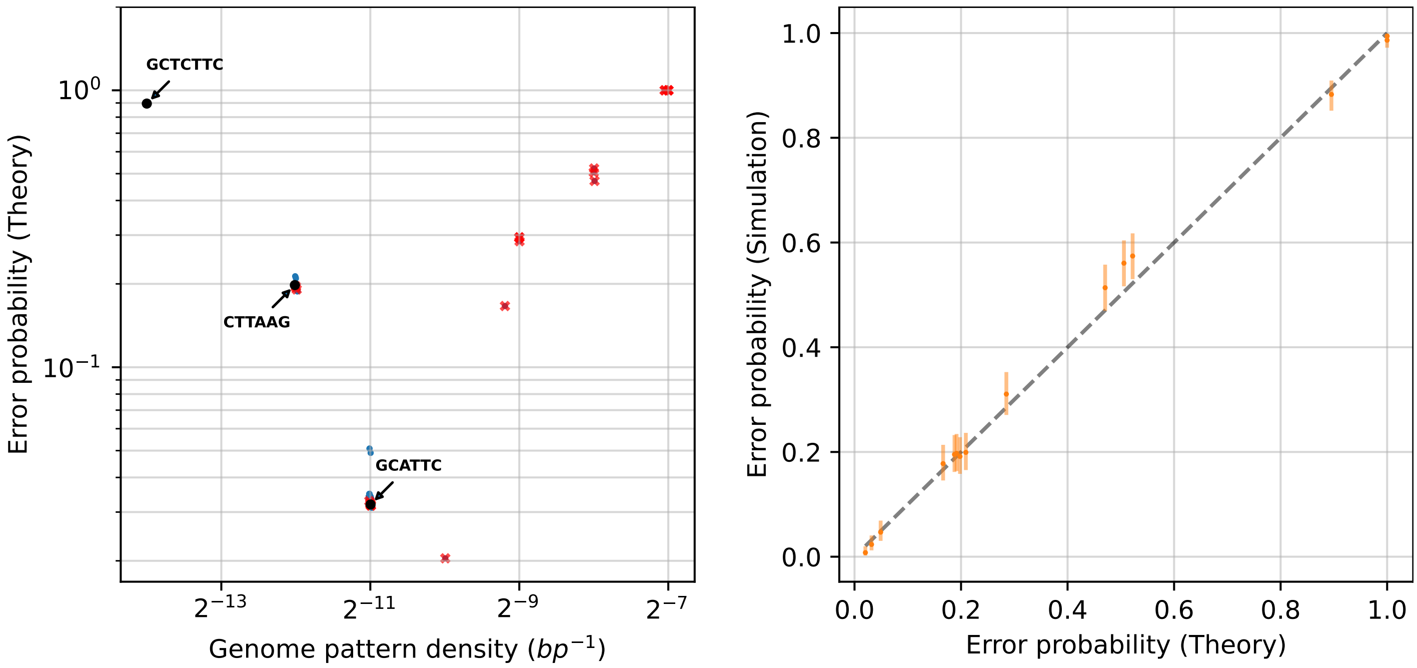


Figure S2: Error probability vs Pattern for a random genome. Here, a random genome sequence of length $10^8$ bp was used. The same patterns as in Figure 4 are shown. As expected for a random sequence, all palindromic 6-letter patterns over $\{A, C, G, T\}$ have an approximate density of $4^{-6}$, while the non-palindromic patterns have twice the density, as both the ordinary and the reverse complement of the pattern are counted.
